# KLF4 induces Mesenchymal - Epithelial Transition (MET) by suppressing multiple EMT-inducing transcription factors

**DOI:** 10.1101/2021.08.26.457621

**Authors:** Ayalur Raghu Subbalakshmi, Sarthak Sahoo, Isabelle McMullen, Aaditya Narayan Saxena, Sudhanva Kalasapura Venugopal, Jason Somarelli, Mohit Kumar Jolly

**Author notes:** Authors to whom correspondence to be addressed (J.A.S), (M.K.J.).

## Abstract

Epithelial-Mesenchymal Plasticity (EMP) refers to reversible dynamic processes where cells can transition from epithelial to mesenchymal (EMT) or from mesenchymal to epithelial (MET) phenotypes. Both these processes are modulated by multiple transcription factors acting in concert. While EMT-inducing transcription factors (TFs) – TWIST1/2, ZEB1/2, SNAIL1/2/3, GSC, FOXC2 – are well-characterized, the MET-inducing TFs are relatively poorly understood (OVOL1/2, GRHL1/2). Here, using mechanism-based mathematical modeling, we show that the transcription factor KLF4 can delay the onset of EMT by suppressing multiple EMT-TFs. Our simulations suggest that KLF4 overexpression can promote phenotypic shift toward a more epithelial state, an observation suggested by negative correlation of KLF4 with EMT-TFs and with transcriptomic based EMT scoring metrics in cancer cell lines. We also show that the influence of KLF4 in modulating EMT dynamics can be strengthened by its ability to inhibit cell-state transitions at an epigenetic level. Thus, KLF4 can inhibit EMT through multiple parallel paths and can act as a putative MET-TF. KLF4 associates with patient survival metrics across multiple cancers in a context-specific manner, highlighting the complex association of EMP with patient survival.

## Introduction

Cancer is expected to surpass all non-communicable disease related deaths in the 21^st^ century, making it a major global public health threat [1]. Nearly all cancer-related deaths can be attributed to the process of metastasis [2]. Metastasizing cells possess the ability of migration and invasion, which enables them to break away from the primary tumour [3], but the process has attrition rates as high as >99.5% [4]. Thus, only a miniscule percentage of metastasizing cells comprise the successful seeding of secondary tumour(s). A key hallmark exhibited by these metastasizing cells is their property of phenotypic plasticity, i.e. their ability to dynamically switch between phenotypes, empowering them to adapt to the ever-changing microenvironments with which they are faced during the metastatic process [5,6]. Therefore, it is critical to decode the mechanisms of phenotypic plasticity in order to unravel the dynamics of metastasis and develop therapeutic strategies targeting this currently-insurmountable challenge.

A major type of phenotypic transition during metastasis is epithelial-mesenchymal plasticity (EMP), i.e. the bidirectional switching among epithelial, mesenchymal and one or more hybrid epithelial/mesenchymal (E/M) phenotypes [7]. Many transcription factors (TFs) capable of inducing an Epithelial-Mesenchymal Transition (EMT) are well-characterized, but those driving the reverse of EMT – a Mesenchymal-Epithelial Transition (MET) – remain relatively poorly investigated. For instance, ZEB1/2, SNAI1/2, TWIST and GSC (Goosecoid) are EMT-TFs that are often activated by signaling pathways, such as TGFβ, and can drive varying extents of EMT in cancer cells through repressing various epithelial genes (such as E-cadherin) and/or inducing the expression of mesenchymal genes (such as vimentin) [8–13]. On the other hand, GRHL1/2 and OVOL1/2 are MET-inducing transcription factors (MET-TFs) that often engage in mutually inhibitory feedback loops with EMT-TFs [14–18]. Recent studies have focused on characterizing drivers and stabilizers of hybrid E/M phenotypes [19–23], which have been claimed to be the ‘fittest’ for metastasis due to their higher plasticity and tumor-initiation potential, and ability to drive collective migration [24], manifested as clusters of circulating tumor cells [25] – the primary harbingers of metastasis [26]. The role of hybrid E/M cells in metastasis is supported by clinical studies demonstrating an association of hybrid E/M features with worse clinicopathological traits [27–29]. However, to effectively target the hybrid E/M phenotype(s), a better understanding of the emergent dynamics of various coupled intracellular and intercellular regulatory networks involved in partial and/or full EMT/ MET is required [30].

Krüppel-like factor 4 (KLF4) is an evolutionarily-conserved zinc finger-containing transcription factor [31]. It is associated with terminal differentiation and homeostasis of various epithelial tissues, including its role in maintaining the stability of adherens junctions and establishing barrier function of the skin [32–34]. It also helps maintain proliferative and pluripotency properties of embryonic stem cells [35] and is crucial for somatic cell reprogramming [32]. Recently, KLF4 has been investigated in the context of EMT. For instance, in corneal epithelial homeostasis, KLF4 upregulates the levels of various epithelial markers, such as E-cadherin and claudins, and downregulates mesenchymal markers, such as vimentin and nuclear localization of β-catenin [36]. KLF4 inhibits EMT in corneal epithelium by preventing phosphorylation and nuclear localization of SMAD2, thus attenuating TGF-β signaling [37]. Similarly, in pulmonary fibrosis, KLF4 inhibits TGFβ1-induced EMT in human alveolar epithelial cells [38]. In tumor progression, it has been proposed as both an oncogene and as a tumor suppressor, depending on the context [39–42]. Thus, a deeper understanding of the roles of KLF4 in tumor progression is needed.

At a molecular level, KLF4 has been shown to inhibit, and be inhibited by, both SNAIL (*SNAI1*) [43,44] and SLUG (*SNAI2*) [45], two of the members of the *SNAI* superfamily that can induce EMT to varying degrees [9,46]. Such a mutually inhibitory feedback loop (also described as a ‘toggle switch’) has also been reported between a) miR-200 and ZEB [47], b) SLUG and SNAIL [48], and c) SLUG and miR-200 [48]. Thus, KLF4, SNAIL and SLUG form a ‘toggle triad’ [49]. In addition, KLF4 can self-activate [50], similar to ZEB1 [51], while SNAIL inhibits itself and activates ZEB [48].

Here, we developed a mechanism-based mathematical model that captures the abovementioned interactions to decode the effects of KLF4 on EMT. Our model predicts that KLF4 can inhibit the progression of EMT by inhibiting the levels of various EMT-TFs; consequently, its overexpression can induce a partial or complete MET, similar to observations for GRHL2 [52–55]. Analysis of *in vitro* transcriptomic datasets and cancer patient samples from The Cancer Genome Atlas (TCGA) reveals a negative correlation between KLF4 levels and EMT score. We also incorporate the impact of epigenetic influence mediated by KLF4 and SNAIL in a population dynamics scenario, and demonstrate that KLF4-mediated ‘epigenetic locking’ can enable resistance to EMT, while SNAIL-mediated effects can drive a stronger EMT. Together, we propose KLF4 as a potential MET-TF that can repress many EMT-TFs simultaneously and inhibit EMT through multiple parallel paths. These observations are supported by the observed association of KLF4 with patient survival metrics across multiple cancers.

## Results

### KLF4 inhibits the progression of EMT

We began with examining the role of KLF4 in modulating EMT dynamics. We investigated the dynamics of interaction between KLF4 and a core EMT regulatory circuit (denoted by the black dotted rectangle in **Fig 1A**) comprised of four players: three EMT-inducing transcription factors (EMT-TFs) - ZEB, SNAIL, SLUG - and an EMT-inhibiting microRNA family (miR-200).

**Fig 1:**
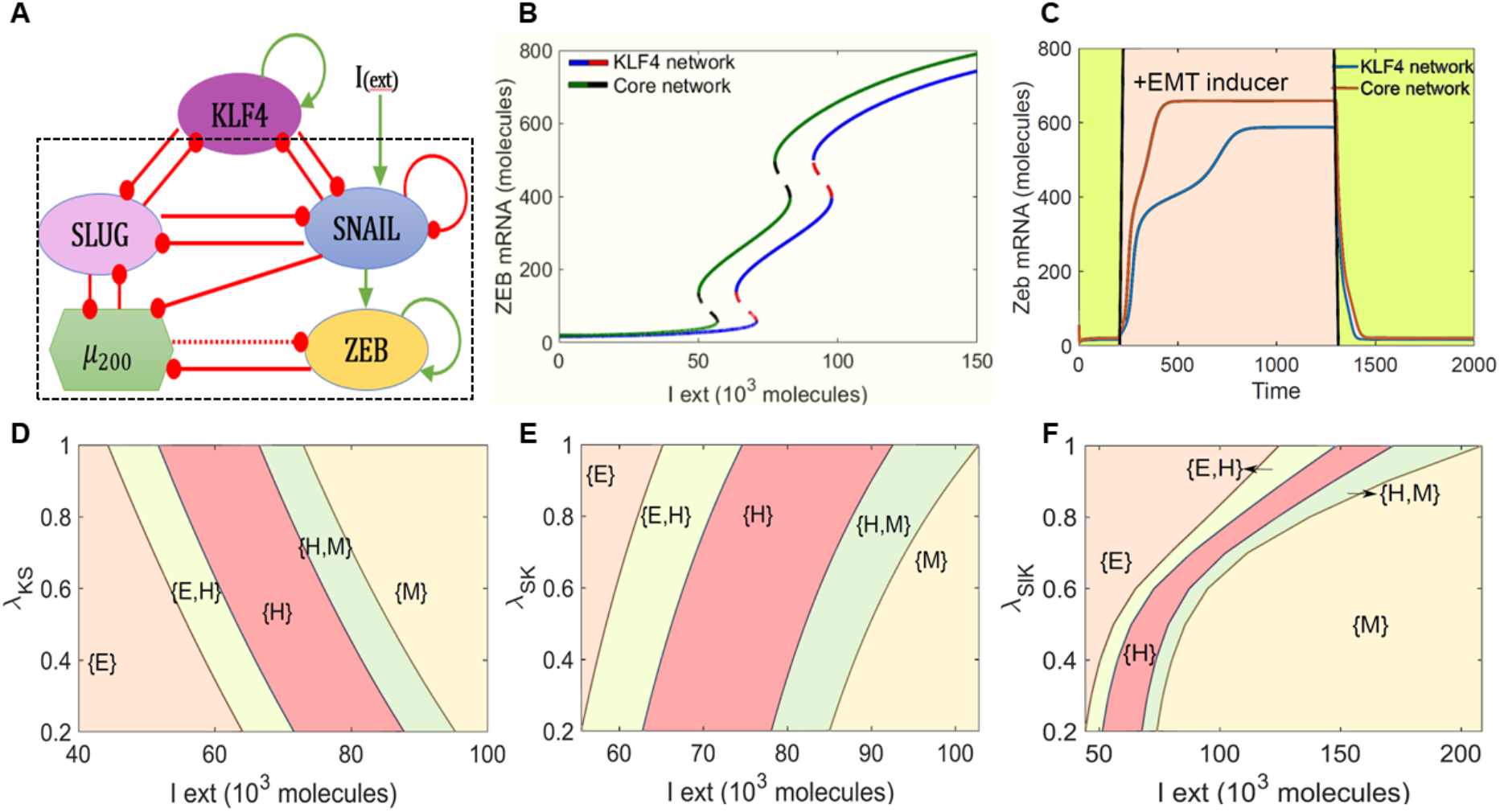
KLF4 inhibits EMT. **A)** Schematic representation of KLF4 coupled to EMT regulatory network consisting of miR-200, ZEB1, SNAIL and SLUG. Green arrows denote activation and red bars indicate inhibition. Solid arrows represent transcriptional regulation; dotted line represents micro-RNA mediated regulation. Circuit shown within the dotted rectangle is the control case (i.e. core EMT network without KLF4). **B)** Bifurcation diagrams indicating ZEB mRNA levels for increasing external signal (I) levels for the coupled EMT-KLF4 circuit (solid blue and dotted red curve) and the core EMT circuit (solid green and dotted black curve). **C)** Temporal dynamics of ZEB mRNA levels in a cell starting in an epithelial phenotype, when exposed to a high level of external EMT signal (I_ext= 100,000 molecules) (pink-shaded region) for circuits shown in A). **D-F)** Phase diagrams for the KLF4-EMT network driven by an external signal (I_ext) for **D)** varying strength of repression on SNAIL by KLF4, **E)** for varying strength of repression on KLF4 by SNAIL, and **F)** for varying strength of repression on SLUG by KLF4.

First, we calculate a bifurcation diagram of ZEB mRNA levels for an external EMT-inducing signal I_ext_, which can represent various possible intracellular/extracellular stimuli that can drive an EMT (**Fig 1B**). ZEB mRNA levels serve as a readout of EMT phenotypes. We observed that with an increase in the strength of I_ext_, the cells can switch from an epithelial state (ZEB mRNA < 100 molecules) to a hybrid E/M phenotype (100 < ZEB mRNA molecules < 600) and finally to a mesenchymal state (ZEB mRNA > 600 molecules). This behaviour is observed both in the absence of KLF4 (curve with green solid line and black dashed line in **Fig 1B**) and presence of KLF4 (curve with blue solid line and red dashed line in **Fig 1B**) and the bifurcation curves look quite similar in shape. However, in the presence of KLF4, we observed that the system required a stronger EMT signal to be pushed out of an epithelial state and also for the acquisition of a completely mesenchymal state (i.e. the bifurcation diagram in the presence of KLF4 is shifted to the right as compared to the core network). This observation suggests that KLF4 can inhibit the progression of EMT.

To test the robustness of the model prediction for the role of KLF4 in EMT, we performed sensitivity analysis in which we varied the numerical value of every kinetic parameter used in the model by ± 10% one at a time, and captured the change in the range of the I_ext_ for which the hybrid E/M state existed in the bifurcation diagram. Except a few parameter cases involving ZEB and miR200 interactions, this change was found to be less than 10% for a corresponding 10% change in the parameter values. Specifically, for variation in parameters indicating the interactions of KLF4 with the core EMT circuit, this change did not extend beyond 1% (**Fig S1A**). Thus, the observed behavior of KLF4 in its ability to delay or inhibit EMT looks to be robust to small parametric variations.

Next, we determined the temporal response of the cell to a fixed concentration of the external EMT-inducing signal I_ext_. In the absence of KLF4, a cell in the epithelial state transitioned first to a hybrid E/M state and then to a mesenchymal state in response to an external signal (red curve in Fig **1C**). But in the presence of KLF4, this transition was much more gradual and relatively slower (blue curve in **Fig 1C**). In addition, the steady state value of ZEB mRNA levels was lower in the presence of KLF4 as compared to the control case. This decrease can be attributed to KLF4-mediated inhibition of both SLUG and SNAIL that can activate ZEB and is consistent with trends in the ZEB mRNA level bifurcation diagram (the blue curve lies below the green curve at all values of I_ext_ in **Fig 1B**).

KLF4 inhibits both SLUG and SNAIL and is inhibited by both of them. Thus, we probed the impact of interactions between KLF4 and both of these EMT-TFs in terms of influencing EMT progression. First, we varied the strength of repression of SNAIL by KLF4. When the strength of this repression was high (i.e., low λ_KS_ value or low K^0^S value), the cells required a stronger EMT-inducing signal to be pushed out of the epithelial state. Conversely, as the value of λ_KS_ or K^0^S increased (i.e., KLF4 inhibits SNAIL weakly), EMT can be induced at lower values of I_ext_ (**Fig 1D, S1B**). Next, we varied the repression of KLF4 by SNAIL. At stronger repression (i.e., low λ_SK_ value or low S^0^K value), cells could exit the epithelial state at a weaker external EMT-inducing signal. Conversely, as the value of λ_SK_ or S^0^K increased (i.e., SNAIL inhibits KLF4 weakly), a stronger stimulus was required for cells to exit the epithelial state (**Fig 1E, S1C**). Put together, these results highlight that while a weaker impact of KLF4 – through either stronger repression of KLF4 by SNAIL or by weaker repression of SNAIL by KLF4 – potentiated the progression of EMT, a stronger impact of KLF4 attempted to prevent cells from undergoing EMT. Similar results were seen for the feedback loop between SLUG and KLF4 (**Fig 1F, S1D-E**), but the impact on EMT dynamics was weaker upon altering the inhibition of SLUG by KLF4 than that of SNAIL by KLF4. Upon altering either λ_KSl_ or K^0^Sl, we did not observe any change in the concentration of I_ext_ needed to induce EMT, as seen for the case with SNAIL (compare **FigS1D** with **Fig1D**, and **Fig S1E** with **Fig S1B**). This difference may be explained by reports suggesting that SNAIL is a more potent EMT inducer than SLUG [9,46]. This hypothesis is strengthened by observations that SLUG self-activation does not alter the qualitative dynamics of KLF4-SNAIL interactions (**Fig S2A-C**). A stronger KLF4 self-activation, on the other hand, can increase the resistance to undergo EMT (**Fig S2D**).

### KLF4 promotes an epithelial phenotype

We next examined whether the impact of KLF4 in inhibiting EMT can be a generic emergent property of the topology of the regulatory network that it forms with SLUG, SNAIL and ZEB1, instead of the behavior of a specific parameter set. Thus, to map the possible phenotypic space of the network shown in **Fig1A**, we simulated its dynamics using the computational framework RACIPE [56]. RACIPE takes in the topology of the GRN as an input and converts that network topology information into a set of coupled ordinary differential equations (ODEs). These ODEs are solved over a wide range of biologically relevant parameter values to identify different possible steady state gene expression levels (phenotypes) that the network is capable of giving rise to.

After obtaining possible phenotypes from RACIPE analysis, we plotted the distributions of steady-state levels of different nodes in the GRN obtained across the ensemble of parameter sets. KLF4, SLUG, ZEB1 and miR-200 showed a bimodal distribution, while SNAIL showed a unimodal one (**Fig S3A**); thus, SNAIL was excluded from subsequent clustering analysis. We plotted the steady states obtained from RACIPE derived as a heatmap (**Fig 2A**, left). Hierarchical clustering performed on the heatmap revealed two predominant clusters – one with (high miR200, high KLF4, low ZEB, low SLUG) levels and other with (low miR200, low KLF4, high ZEB, high SLUG) levels. These clusters can be mapped on to epithelial and mesenchymal phenotypes, respectively (green and orange bars in **Fig 2A**). Next, we perturbed the production rates of KLF4 to mimic over- and under-expression of KLF4 and assessed its effect of the frequency of the observed epithelial and mesenchymal phenotypes. Overexpression of KLF4 led to increased frequency of the epithelial cluster, and a decrease in the mesenchymal cluster (**Fig 2A**, right). As expected, opposite patterns were observed upon inhibition and over-expression of ZEB1, an EMT-TF (**Fig S3B**). We quantified the change in the fraction of the epithelial phenotype when KLF4, ZEB1 and SLUG are either overexpressed or inhibited, one at a time. Overexpression of KLF4 or downregulation of either SLUG or ZEB1 increased the frequency of the epithelial phenotype (**Fig 2B**).

**Fig 2:**
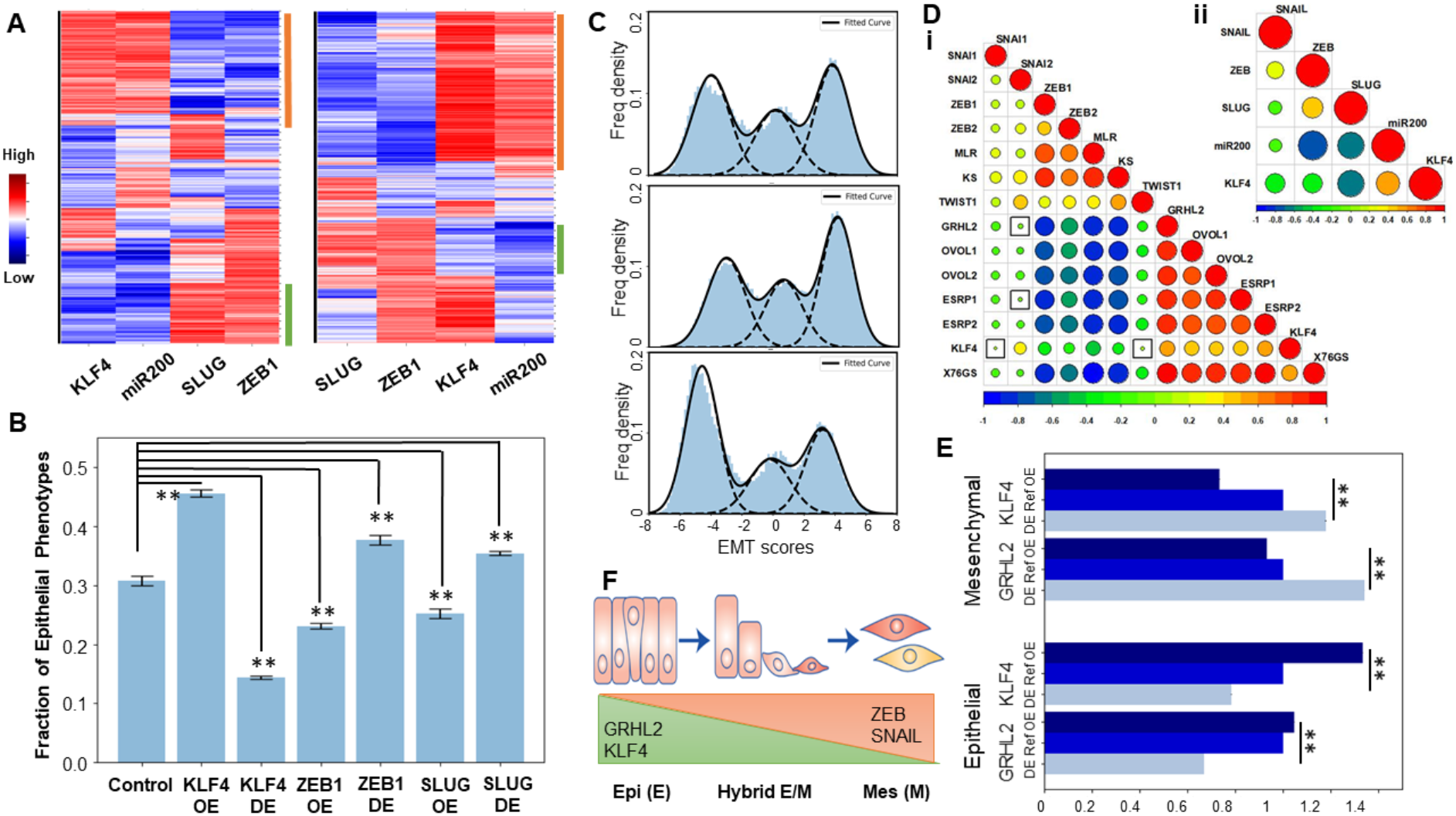
KLF4 promotes an epithelial phenotype. **A)** (Left): Heatmap showing the steady state values of all components of the KLF4-EMT coupled circuit, obtained across parameter sets simulated by RACIPE. (Right) Same as the left panel, but for KLF4 overexpression. **B)** Change in fraction of epithelial phenotype due to simulated overexpression or downregulation of either KLF4 or ZEB1 or SLUG. Error bars for n=3 independent simulations. **C)** Frequency density of the epithelial, hybrid and mesenchymal phenotypes obtained using EMT scores. (Top) KLF4 circuit, (middle): KLF4 circuit with KLF4 down expression (DE), (bottom) KLF4 circuit with KLF4 overexpression (OE). **D)** Pairwise Pearson’s correlation matrix (i) Correlation of epithelial and mesenchymal players and EMT scoring metrics (76GS, KS, MLR) in CCLE cell lines (ii) Correlation of RACIPE simulated expression values of nodes in KLF4 network. Squares indicate p-value>0.01. **E)** Change in size of epithelial and mesenchymal clusters upon OE and DE of GRHL2 and KLF4. **F)** Schematic showing the variation of KLF4, GRHL2, ZEB1 and SNAIL levels across the EMT spectrum. *: p<0.05; **: p <0.01

To investigate the classification of cell phenotypes in a more quantitative manner and to characterize heterogeneity in EMT in a given cell population, we defined an EMT score (= ZEB1 + SLUG – miR-200 – CDH1) for an extended regulatory network that included E-cadherin and its connections with SLUG and ZEB1 (**Fig S3C**). Distribution of EMT scores plotted for solutions seen across parameter sets in RACIPE reveal a trimodal distribution, indicating the existence of three distinct EMT phenotypes (Fig **2C**, top panel). Simulated knockdown of KLF4 drove an increase in mesenchymal subpopulation with a concurrent decrease in epithelial states (**Fig 2C**, middle panel). Opposite trends were observed for case of simulated KLF4 over-expression (**Fig 2C**, bottom panel). These results highlight that the emergent dynamics of the KLF4-EMT network can allow for multiple phenotypes to co-exist in an isogenic population; and KLF4 levels can modulate the distribution of phenotypic heterogeneity in that population towards a more epithelial or a more mesenchymal state.

To interrogate the role of KLF4 in EMT/MET further, we performed pairwise correlation analysis in the Cancer Cell Line Encyclopaedia (CCLE) cohort among three transcriptomic-based EMT scoring metrics (76GS, KS, MLR) and expression levels of KLF4 and those of other canonical epithelial and mesenchymal factors. KLF4 correlated negatively with the KS and MLR EMT scoring metrics (higher KS or MLR scores denote a mesenchymal phenotype [57]) but positively with 76GS scores (higher 76GS scores denote a more epithelial phenotype [57]) (**Fig 2D**, i). Most EMT-TFs were found to be correlated positively with each other (SNAI1/2, ZEB1/2, TWIST1) and negatively with KLF4 and other MET drivers, such as ESRP1/2, OVOL1/2 and GRHL2 [58], which were all positively corelated with KLF4 (**Fig 2D**, i). Consistent correlations were recapitulated in RACIPE simulation data for the KLF4-EMT network (**Fig 2D**, ii), thus underscoring that the GRN considered in Fig 1A can explain these observed experimental trends for the existence of “teams” [59] of EMT and MET inducers. Interestingly, GRHL2 seemed to correlate more strongly with ZEB1, ZEB2, TWIST1 and the MLR and KS scores as compared to KLF4 (**Fig 2D**, i), thus encouraging us to compare the influence of KLF4 and GRHL2 in terms of their ability to induce MET via simulations. We compared the OE and DE scenarios of GRHL2 and KLF4 in terms of influencing the distribution of epithelial and mesenchymal phenotypes, and noted a stronger enrichment of mesenchymal upon down-regulation of GRHL2 than that seen upon down-regulation of KLF4 (**Fig 2E; S3D**). Thus, our results suggest that KLF4, similar to GRHL2, can induce a partial or full MET (**Fig 2F**).

### KLF4 is inhibited during EMT

Next, using various publicly available transcriptomic datasets, we examined if KLF4 is inhibited as cells undergo EMT. In mouse mammary cells EpRas induced to undergo EMT by TGFβ treatment for 14 days [60], KLF4 levels were reduced (GSE59922; **Fig 3A**). Similarly, when EMT was induced in HMEC cells via overexpression of SNAIL or SLUG [9], KLF4 levels went down (GSE40690; **Fig 3B**). Reinforcing trends were seen in MCF-7 cells forced to undergo EMT via overexpression of SNAIL [61] (GSE58252; **Fig 3C**), in OVCA4209 cells in which GRHL2 was knocked down [62] (GSE118407; **Fig 3D**), and in MCF10A cells cultured without growth factors and shown to undergo EMT [63] (GSE85857; **Fig 3E**). Further, as compared to the primary HMEC (human mammary epithelial cells), immortalized and Ras-transformed HMECs were enriched for EMT-associated genes (GSE110677) and had lower KLF4 levels (**Fig 3F**). Together, these datasets across cancer types reveal a robust reduction in KLF4 expression with the onset of EMT.

**Fig 3:**
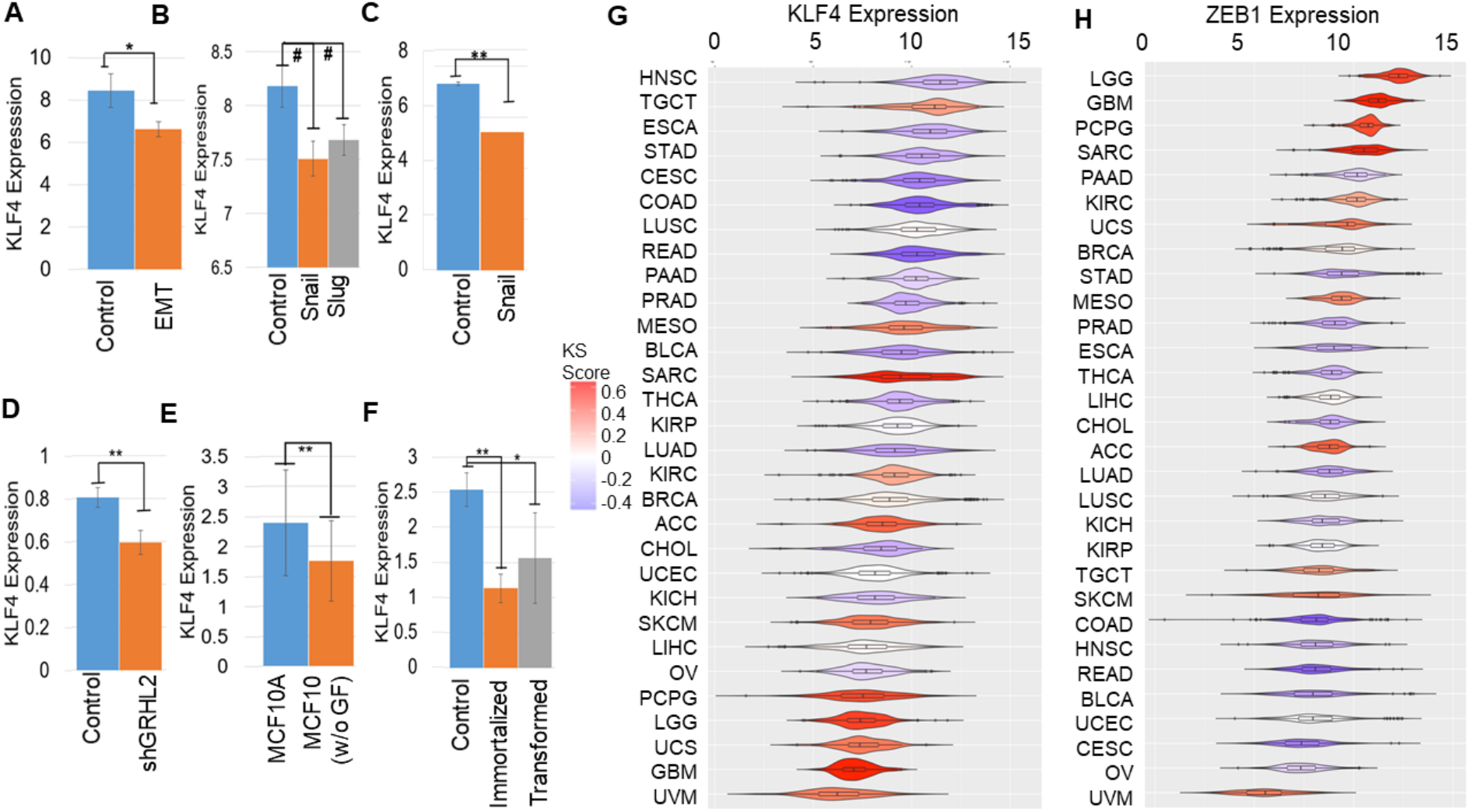
KLF4 is inhibited during EMT. KLF4 expression across comparison groups in GEO datasets. **A)** GSE59922 **B)** GSE40690 **C)** GSE58252 **D)** GSE118407 **E)** GSE85857 **F)** GSE110677. #: p <0.1, *: p< 0.05; **: p<0.01 for Student’s two-tailed t-text with unequal variances. **G, H)** KLF4 and ZEB1 expression in TCGA cancer types, arranged by mean KS scores (color scheme given on the right).

To substantiate these *in vitro* observations with clinical data, we compared KLF4 levels across cancers in The Cancer Genome Atlas (TCGA). KLF4 expression was found to be lower in cancers with a more mesenchymal phenotype as measured by their higher KS-based EMT scores [64]. Mesenchymal-like cancers, such as uveal melanoma (UVM), uterine carcinosarcoma (UCS), glioblastoma (GBM) and low grade glioma (LGG), tended to have lower KLF4 expression (shown in red in **Fig 3G**). Conversely, KLF4 levels were higher in epithelial-like cancer types, such as head and neck squamous cell carcinoma (HNSC), oesophageal carcinoma (ESCA), stomach adenocarcinoma (STAD) and cervical carcinoma (CESC) samples (blue EMT scores; **Fig 3G**). ZEB1 and SNAIL, on the other hand, showed opposite trends to KLF4: enriched in cancers with a higher KS score: LGG, GBM, UCS, SARC (sarcoma) and PCPG (pheochromocytoma and paraganglioma), but reduced in those with a lower one: HNSC, COAD (colorectal adeno-carcinoma), CESC, BLCA (bladder carcinoma) and READ (rectum adenocarcinoma) (**Fig 3H, S4A**). Hence, an inverse correlation of KLF4 with multiple EMT-TFs seen *in vitro* is consistently observed in TCGA samples.

### Epigenetic changes, including KLF4 promoter methylation, can alter population distributions along the EMT spectrum

Decreased KLF4 expression has been reported to be associated with hypermethylation of the KLF4 promoter during EMT in renal fibrosis *in vitro* and *in vivo* [65]. Thus, we examined the correlation of KLF4 expression with its methylation status in TCGA data. We observed reduced methylation of KLF4 in many cancers with reduced KS scores such as HNSC, ESCA, COAD. Consistent with this observation, KLF4 expression and methylation status were negatively correlated (**Fig 4A**), reminiscent of observations in renal cancer cell lines and tissues, and suggesting a possible epigenetic mechanism driving its suppression during EMT. Consistently, DNA methyltransferase inhibitor increased KLF4 expression in renal cancers [66]. SNAIL expression was also negatively correlated with corresponding promoter methylation levels in TCGA; however, ZEB1 did not show a clear pattern (**Fig S4B,C**). These observations drove us to investigate the impact of epigenetic influence operating in the KLF4 and SNAIL feedback loop.

**Fig 4:**
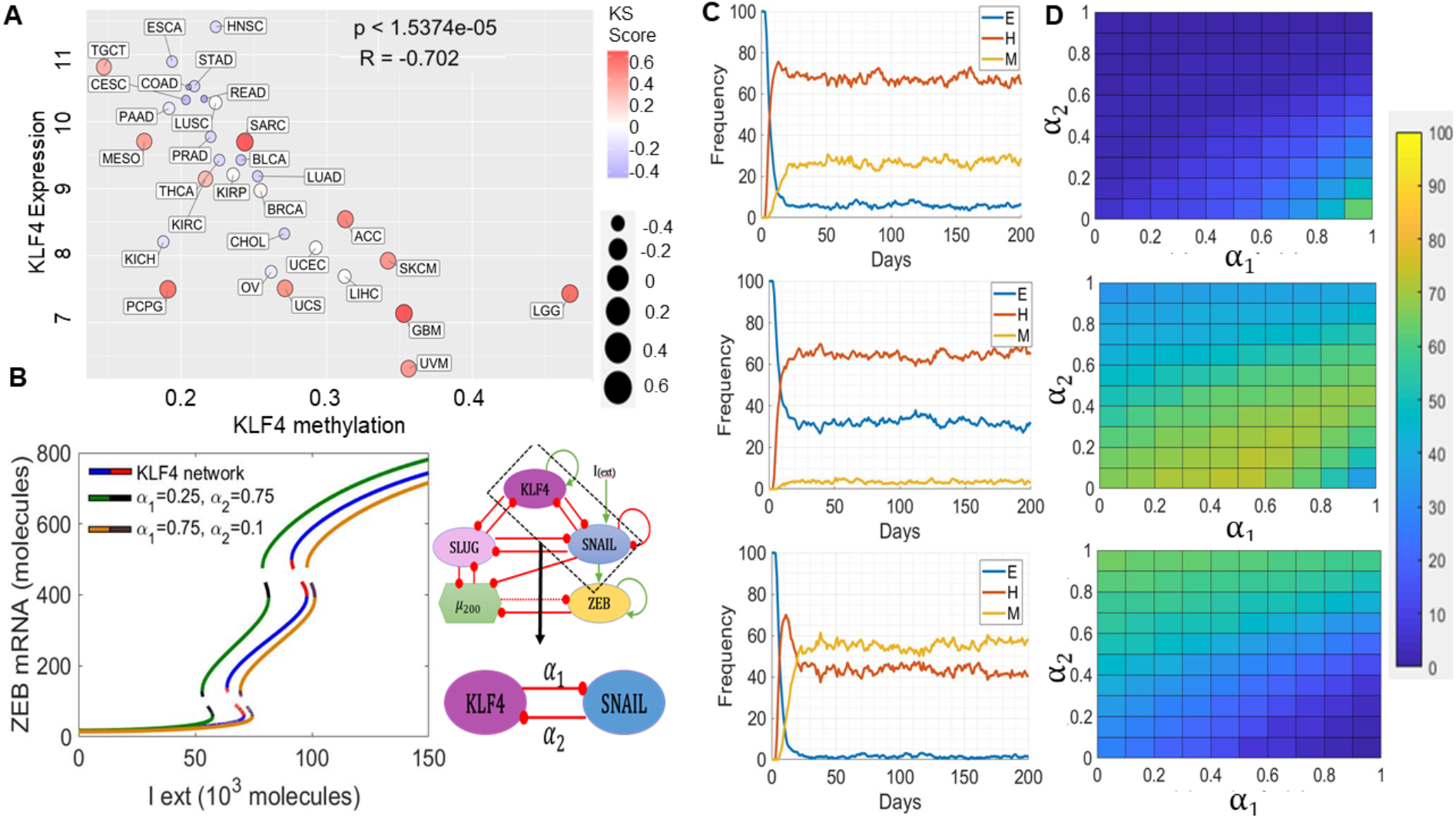
Epigenetic modulations involving KLF4 can alter the population dynamics of EMT states. **A)** Scatter plot for KLF4 expression and its methylation status in TCGA cancer types. **B)** Bifurcation diagrams indicating the ZEB mRNA levels for increasing EMT-inducing external signal (I_ext) levels for the coupled EMT-KLF4 circuit (solid blue and dotted red curve), for the circuit with higher α_1_ and lower α_2_ values (solid yellow and dotted brown curve) and for the circuit with lower α_1_ and higher α_2_ values (solid green and dotted black curve). **C)** Stochastic simulation of KLF4-EMT network for varied values of α_1_ and α_2_. (Top) α_1=_ α_2_=0, (middle) α_1_=0.75 α_2_=0.1, (bottom) α_1_=0.25 α_2_=0.75. **D)** Population distribution of E (top), hybrid E/M (middle) and M (bottom) cells for varying values of α_1_ and α_2_.

Epigenetic changes can drastically alter the rates of transition among cell phenotypes by controlling the accessibility of promoters for ‘master regulators’. In the context of EMT, we have previously shown that epigenetic feedback mediated by ZEB1 while repressing miR-200 (i.e. blocking access to miR-200 promoter for MET inducers) can drive irreversible EMT, while that mediated by GRHL2 (i.e. blocking access to ZEB1 promoter for EMT inducers) in inhibiting ZEB1 can enable irreversible MET, i.e. resistance of cells to undergo EMT [67,68]. Here, we assessed the impact of KLF4-mediated epigenetic silencing of SNAIL (i.e. the ability of KLF4 to cause methylation of SNAIL promoter directly or indirectly) and *vice versa* (SNAIL-mediated epigenetic silencing of KLF4) with a population dynamics model capturing a cell population with diverse EMT states (epithelial, mesenchymal and hybrid E/M). This phenomenological model encapsulates epigenetic influence by modulating the threshold for the impact of a transcription factor on expression of its downstream target [69]. Such dynamic thresholds capturing epigenetic influence often enable self-stabilization of gene expression states, i.e. the longer a transcription factor has been active, the easier it becomes for it to stay “on”. We introduced two epigenetic variables: α1 and α2. The higher the value of α1, the stronger the influence of KLF4-mediated effective epigenetic silencing of SNAIL. The higher the value of α2, the stronger the influence of SNAIL-mediated effective epigenetic silencing of KLF4 (see **Methods** for details).

As a first step towards understanding the dynamics of this epigenetic ‘tug of war’ between KLF4 and SNAIL, we characterized how the bifurcation diagram of KLF4-EMT coupled circuit changed at various values of (α1, α2). When the epigenetic silencing of SNAIL was higher in the context of KLF4 expression ((α1, α2) = (0.75, 0.1)), a larger EMT-inducing signal (I_ext) is required to push cells out of an epithelial state, because SNAIL is being strongly repressed by KLF4 as compared to the control (no epigenetic influence) case (compare the blue/red curve with the black/yellow curve in **Fig 4B**). Conversely, when the epigenetic silencing of KLF4 predominated in the context of SNAIL expression ((α1, α2) = (0.25, 0.75)), it is easier for cells to exit an epithelial state, because the KLF4 repression of EMT is now being inhibited more potently by SNAIL relative to the control case (compare the blue/red curve with the black/green curve in **Fig 4B**). Thus, these opposing epigenetic “forces” can “push” the bifurcation diagram in different directions along the x-axis, without impacting any of its major qualitative features.

To consolidate these results, we next performed stochastic simulations for a population of 500 cells at a fixed value of I_ext = 90,000 molecules. We observed a stable phenotypic distribution with 6% epithelial (E), 28% mesenchymal (M) and 66% hybrid E/M cells (**Fig 4C**, top) in the absence of any epigenetic regulation (α1= α2 = 0). In the case of a stronger epigenetic repression of SNAIL by KLF4 (α1 =0.75, α2 = 0.1), the population distribution changed to 32% epithelial (E), 3% mesenchymal (M) and 65% hybrid E/M cells (**Fig 4C**, middle). Conversely, when SNAIL repressed KLF4 more dominantly (α1 =0.25, α2 = 0.75), the population distribution changed to 2% epithelial (E), 58% mesenchymal (M) and 41% hybrid E/M cells (Fig **4C**, bottom). A similar analysis was performed for collating steady-state distributions for a range of α1 and α2 values, revealing that high α1 and low α2 values favoured the predominance of an epithelial phenotype (Fig **4F**, top), but low α1 and high α2 values facilitated a mesenchymal phenotype (Fig 4F, bottom). Intriguingly, when the strength of epigenetic repression from KLF4 to SNAIL and *vice versa* was comparable, the hybrid E/M phenotype dominated (**Fig 4F**, middle). Put together, varying extents of epigenetic silencing mediate by an EMT-TF SNAIL and a MET-TF KLF4 can fine-tune the epithelial-hybrid-mesenchymal heterogeneity patterns in a cell population.

### KLF4 correlates with patient survival

To determine the effects of KLF4 on clinical outcomes, we investigated the correlation between KLF4 and patient survival. We observed that high KLF4 levels correlated with better relapse-free survival (**Fig 5A-B**) and better overall survival (**Fig 5C-D**) in breast cancer samples. We also examined the overall survival data relative to KLF4 expression in additional clinical datasets across lung cancer and ovarian cancer. KLF4 associated with overall survival in a context-specific manner, with some data sets showing an improved overall survival with high KLF4 and others the opposite trend (**Fig 5E**).

**Fig 5:**
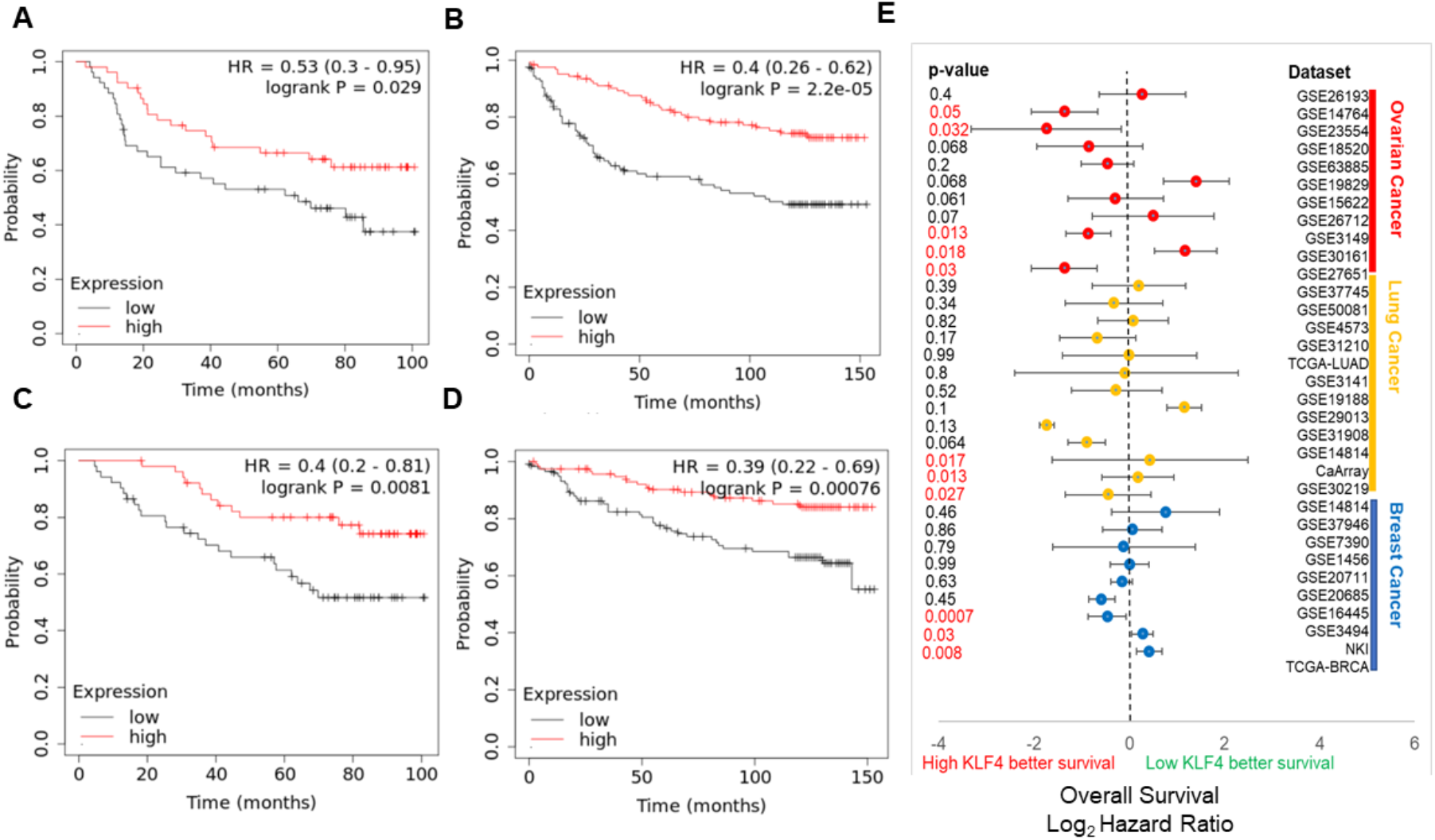
KLF4 correlates with patient survival in a context-specific manner. **A,B)** Relapse free survival trends in GSE42568 and GSE3494 (breast cancer), respectively. **C,D)** Overall survival trends in GSE42568 and GSE3494 (breast cancer), respectively. HR denotes hazard ratio; PVAL denotes p-value. **E)** Plot of log_2_ hazard ratio for overall survival in different cancer types comparing high and low KLF4 expression levels. Mean value and 95% confidence interval are shown.

## Discussion

We hereby propose KLF4 as a potential MET-inducing transcription factor (MET-TFs) based on *in silico* model predictions and their experimental validation across multiple *in vitro* and cancer patient sample data sets. This observation adds to the increasing literature on the role of KLF4 in inhibiting EMT and/or driving MET in different biological contexts. For instance, in colon epithelial cell line RKO, KLF4 upregulates the levels of various epithelial-specific keratins, such as KRT8 and KRT18 [70]. Similarly, in nasopharyngeal carcinoma, KLF4 can transcriptionally activate E-cadherin and reduce the motility and invasion of cells. This reduction is at least partly rescued by shRNA-mediated E-cadherin knockdown in KLF4-expressing cells, suggesting a functional role of E-cadherin in regulating these traits [71]. Direct transcriptional activation of CDH1 (E-cadherin) by KLF4 has also been noted in MCF10A cells [72]. Further, overexpression of KLF4 in MDA-MB-231 breast cancer cells restored E-cadherin levels, induced an epithelial morphology, and suppressed migration and invasion [72], similar to previous observations in these cells for another MET-TF, GRHL2 [73]. Consistently, KLF4 over-expression decreased the levels of vimentin and Slug in and increased those of E-cadherin in OVCAR3 ovarian cancer cells [74]. These observations are reminiscent of the effect of KLF4 knockdown in a prostate stem cell line where the cells lost their epithelial markers, such as E-cadherin, ZO-1 and cytokeratin 8, and showed elevated levels of Vimentin, SNAIL, SLUG and ZEB1 [75]. Supporting these *in vitro* observations, in pancreatic cancer samples, KLF4 correlated positively with E-cadherin and negatively with vimentin and Cav-1, a direct transcriptional target of KLF4 that can inhibit EMT in pancreatic cancer [76].

KLF4 can also promote stemness in various cancers where it promotes epithelial differentiation, thereby challenging the tacitly assumed association between EMT and cancer stem cells (CSCs) [77]. In breast cancer, KLF4 knockdown reduced the ALDH1+ CSCs and mammosphere formation *in vitro* in MCF7 and MDA-MB-231 cells [41]. *In vivo* tumorigenesis and metastasis was also compromised in KLF4-depleted NOD/SCID mice [41,78]. In hepatocellular carcinoma, KLF4 directly activated EpCAM, increased the number of EpCAM+/CD133+ liver cancer stem cells *in vitro* and amplified tumorigenesis *in vivo* [79]. Similarly, in osteosarcoma cells, KLF4 suppression prevented sphere formation and attenuated the levels of various stem cell-related markers, including ALDH1A1 [80]. Conversely, KLF4 overexpressing cells were more chemoresistant and metastatic [81], and osteosarcoma stem cells have increased levels of KLF4 [82].

However, the association of KLF4 with metastasis is relatively ambiguous and context-dependent. High KLF4 was shown to prevent metastasis in breast cancer [83] and pancreatic cancer models [84]. KLF4 can have additional roles in modulating metastasis beyond regulating EMT and/or stemness, such as in its pro-metastasis role in the phenotypic switching of perivascular cells and formation of pre-metastatic niche [85]. Another potential confounding factor is the association of KLF4 with cell cycle regulation. KLF4 can drive cell cycle arrest [70], but also simultaneously repress p53 and activate p21^CIP^ [42] acting as a ‘incoherent signal’ that can drive antagonistic outputs depending on relative strengths of regulation. Therefore, future studies investigating the coupled dynamics of EMT, stemness and cell cycle and the connection of KLF4 to these regulatory pathways are needed to elucidate the impact of KLF4 in modulating these interconnected axes driving metastasis.

Overall, our analysis highlights KLF4 as a potential MET-TF, adding to the list of known MET-TFs such as GRHL1/2/3 and OVOL1/2. Recent studies showing that MET is not simply the inverse of EMT [12,86,87] necessitate attention to identify more MET-TFs and the interconnected networks they form with EMT-TFs to gain a comprehensive understanding of the emergent dynamics of EMP in the metastatic-invasion cascade.

## Materials and Methods

### Mathematical Modelling

As per the schematic shown in Figure 1A, the dynamics of all the four molecular species (miR-200, SNAIL, ZEB and SLUG) were described by a system of coupled ODEs. The generic chemical rate equation given below describes the level of a protein, mRNA or micro-RNA (X):

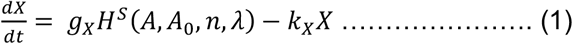

where the first term g_X_ signifies the basal rate of production; the terms multiplied to g_X_ represent transcriptional/translational/post-translational regulations due to interactions among the species in the system, as defined by the shifted Hills function (*H*^*S*^(*A, A*_0_, *n, λ*)). The rate of degradation of the species (X) is defined by the term k_X_X based on first order kinetics. The complete set of equations and parameters are presented in the Supplementary Material. Bifurcation diagrams were drawn in MATLAB (MathWorks Inc.) using the continuation software package MATCONT [88].

### RACIPE (random network simulation)

The gene regulatory network shown in **Fig 1A** was simulated via RACIPE. Over expression and down expression of KLF4, ZEB1 and SLUG was done by setting fold change value to 10. 10000 parameter sets were simulated for 100 different initial conditions to obtain the ensemble of steady state solutions. Steady state solutions were Z-normalised for each gene over all steady state values as (observed steady state expression – mean steady state expression) / standard deviation of steady state expression. The resultant normalised steady state solutions were plotted as a heatmap. Significance in difference between distinct groups were accessed by performing Students t-test on three replicates of 10000 parameter sets each.

Next, we incorporated CDH1 to the circuit in **Fig 1A** and simulated the GRN by RACIPE. A similar circuit was also simulated by incorporating GRHL2 but without KLF4. Along with the base circuits, over-expression and down-expression was also done for KLF4 and GRHL2 by 50-fold in their respective circuits. RACIPE steady states were z-normalised as above and EMT score for each steady state was calculated as ZEB1 + SLUG – miR-200 – CDH1. The resultant trimodal distribution was quantified by fitting 3 gaussians. The frequencies of epithelial and mesenchymal phenotypes were quantified by computing the area under the corresponding gaussian fits. Significance in difference between distinct groups were accessed by performing Students’ t-test on three replicates of 10,000 parameter sets each.

### Gene expression datasets

Gene expression datasets were downloaded using the GEOquery R Bioconductor package [89]. Pre-processing of these datasets was performed for each sample to obtain the gene-wise expression from probe-wise expression matrix using R (version 4.0.0).

### External signal noise and Epigenetic feedback on KLF4 and SNAIL

The external noise and epigenetic feedback calculations were performed as described earlier [68]

a. <u>Noise on External signal:</u> The external signal I that we use here can be written as the stochastic differential equation:

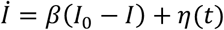

Where, *η*(*t*) satisfies the condition *η*(*t*), *n*(*t*′) ≥ *Nδ*(*t* − *t*′). Here I_0_ is set at 90 K molecules, β as 0.04 hour-1, and N as 80 (K molecules/hour)^2^.
b. Epigenetic feedback: We tested the epigenetic feedback on the KLF4-SNAIL axis. The dynamic equation of epigenetic feedback on inhibition by KLF4 on SNAIL is:

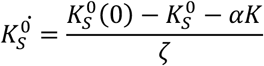

Similarly, epigenetic feedback on SNAIL inhibition on KLF4 is modelled via:

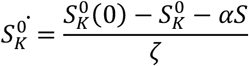

where ζ is a timescale factor and chosen to be 100 (hours). α represents the strength of epigenetic feedback. Larger α corresponds to stronger epigenetic feedback. α has an upper bound because of the restriction that the numbers of all molecules must be positive. For inhibition by KLF4 on SNAIL, high level of KLF4 can inhibit the expression of SNAIL due to this epigenetic regulation. Meanwhile, for SNAIL’s inhibition on KLF4, high levels of SNAIL can suppress the synthesis of KLF4.

### Kaplan-Meier analysis

ProgGene [90] and KM Plotter [91] were used to conduct Kaplan–Meier analysis for respective datasets. The number of samples in KLF4-high vs. KLF4-low categories are given in SI Table.

### Patient Data

Gene expression levels for batch effects normalized RNA-seq were obtained for 12,839 samples from The Cancer Genome Atlas’s (TCGA) pan-cancer (PANCAN) dataset via University of California, Santa Cruz’s Xena Browser. The samples were filtered using unique patient identifiers and only samples that overlapped between the two datasets were kept (11,252 samples). Samples were further filtered to remove patients with missing data for gene expression or cancer type (10,619 samples). These samples were ultimately used in all subsequent analyses. DNA methylation data was obtained from the TCGA PANCAN dataset via University of California, Santa Cruz’s Xena Browser. Methylation data was profiled using Illumina Infinium HumanMethylation450 BeadChip (450K) [92].

#### EMT score

Epithelial-mesenchymal transition (EMT) score was calculated using Kolmogorov– Smirnov (KS) scoring metric [93]. For a given patient, the cumulative distribution functions (CDFs) of Epithelial and Mesenchymal gene signatures were compared. First, the distance between Epithelial and Mesenchymal signatures was calculated using the maximum distance between their cumulative distribution functions. This represents the test statistic in the subsequent two-sample test used to calculate the EMT score. The score was ultimately determined by hypothesis testing of two alternative hypotheses with the null hypothesis being that there was no difference in CDF of Epithelial and Mesenchymal signatures. The first hypothesis was that CDF of Mesenchymal signature is greater than CDF of Epithelial signature. The second hypothesizes that the CDF of Epithelial signatures is greater than the CDF of Mesenchymal signatures. This scoring metric ranges from -1 to +1 where a sample with a positive EMT score is Mesenchymal whereas negative EMT scores are associated with an Epithelial phenotype. MLR and 76GS scores were calculated as earlier [57].

#### Methylation Status

After the four genes of interest were identified as KLF4, SLUG (SNAI2), SNAIL (SNAI1), and ZEB1, methylation and expression data obtained from the TCGA PANCAN dataset via the University of California, Santa Cruz’s Xena Browser. Methylation data was quantified from the Illumina Infinium HumanMethylation450 BeadChip (450K) for each of the four genes and identified unique CpG associated with each of these genes. 10 unique CpG sites for KLF4; 10 for SLUG; 13 for SNAIL; and 42 for ZEB1. A β value, representing how methylated each CpG site was with a β value of one representing a fully methylated CpG site, was provided for all patients and at every CpG site. To quantify the methylation status of each gene, the β values for all associated CpG sites were averaged [92]. Gene expression was then visualized by using R’s ggplot2 package to display violin plots for each gene that were ordered by gene expression and colored by EMT score. Subsequently, the methylation data and expression data were again plotted using R’s ggplot2 package with labels created via R’s ggrepel package.

## Acknowledgements

This work was supported by Ramanujan Fellowship (SB/S2/RJN-049/2018) awarded to MKJ by SERB, Department of Science and Technology, Government of India. JAS is supported by the Department of Defense (W81XWH-18-1-0189) and NCI 1R01CA233585-03. The results published here are based, in part, upon data generated by the TCGA Research Network: https://www.cancer.gov/tcga.

## Conflict of Interest

The authors declare no conflict of interest.

## Author contributions

MKJ designed research; MKJ and JAS supervised research; ARS, SS, IM, AS and SKV performed research; all authors analysed data and contributed to manuscript writing.

## Code availability

All codes used in the manuscript are available on GitHub at https://github.com/ARShuba/KLF4

## Notes

### Competing Interest Statement

The authors have declared no competing interest.

