## Supplementary Information for "KLF4 induces Mesenchymal - Epithelial Transition (MET) by suppressing multiple EMT-inducing transcription factors"

The dynamics of the molecular species of the EMT regulatory circuit (miR-200, Snail, Zeb, Slug) and KLF4 (shown in Fig 1A) is described using coupled ordinary differential equations.

$$\frac{d\mu_{200}}{dt} = g_{\mu_{200}} H^s(Z,\lambda_{Z,\mu_{200}}) H^s(S,\lambda_{S,\mu_{200}}) H^s(Sl,\lambda_{Sl,\mu_{200}}) - m_Z Y_\mu(\mu_{200}) - m_{Sl} Y_\mu(\mu_{200}) - k_{\mu_{200}}\mu_{200}$$

$$\frac{dm_Z}{dt} = g_{m_Z} H^s(Z,\lambda_{Z,m_Z}) H^s(S,\lambda_{S,m_Z}) - m_Z Y_Z(\mu_{200}) - k_{m_Z} m_Z$$

$$\frac{dZ}{dt} = g_Z m_Z L(\mu_{200}) - k_Z Z$$

$$\frac{dS}{dt} = g_S H^s(I,\lambda_{I,S}) H^s(Sl,\lambda_{Sl,S}) H^s(S,\lambda_{S,S}) H^s(K,\lambda_{K,m_S}) - k_S S$$

$$\frac{dm_{Sl}}{dt} = g_{m_{Sl}} H^s(S,\lambda_{S,m_{Sl}}) H^s(K,\lambda_{K,m_{Sl}}) - m_{Sl} Y_Z(\mu_{200}) - k_{m_{Sl}} m_{Sl}$$

$$\frac{dSl}{dt} = g_{Sl} m_{Sl} L(\mu_{200}) - k_{Sl} Sl$$

$$\frac{dK}{dt} = g_K H^s(K,\lambda_{K,K}) H^s(Sl,\lambda_{Sl,S}) H^s(S,\lambda_{S,S}) - k_S S$$

where $g_X$ is the corresponding innate production rate and $k_X$ is the innate degradation rate.

$m_z L(\mu_{200})$ is the net translation rate, $m_z Y_m(\mu_{200})$ is the total ZEB mRNA active degradation rate and $m_z Y_\mu(\mu_{200})$ is the total miR active degradation rate. $H^S$ is the shifted Hill function, defined as

$H^S(B,\lambda) = H^-(B) + \lambda H^+(B)$ ,

$H^-(B) = 1/[1+(B/B_0)^{nB}]$ ,

$H^+(B) = 1 - H^-(B)$,

λ is the fold change from the basal synthesis rate due to protein B. λ >1 for activators, while λ<1 for inhibitors.

λ<1 for inhibitors.

**Parameter Estimation:**

The model parameters were adopted from previously published literature for the molecular species of the core circuit (I, miR-200, Snail, Zeb, Slug) and KLF4 interactions, given below:

| Parameter | Value | Reference |
|---|---|---|
| $g\mu 200$ (Molecules/Hour) | 2.1K | (Lu *et al.*, 2013) |
| gmZ (Molecules/Hour) | 11 | (Lu *et al.*, 2013) |
| $Z^0\mu_{200}$ (Molecules) | 220K | (Lu *et al.*, 2013) |
| $Z^0 m_Z$ (Molecules) | 25K | (Lu *et al.*, 2013) |
| $n_{Z,\mu_{200}}$ | 3 | (Lu *et al.*, 2013) |
| $n_{Z,m_Z}$ | 2 | (Lu *et al.*, 2013) |
| $n_{\mu_{200}}$ | 6 | (Lu *et al.*, 2013) |
| $n_{S,\mu_{200}}$ | 2 | (Lu *et al.*, 2013) |
| $n_{S,m_Z}$ | 2 | (Lu *et al.*, 2013) |
| $\lambda_{Z,\mu_{200}}$ | 0.1 | (Lu *et al.*, 2013) |

| | | |
|---|---|---|
| $\lambda_{Z,m_Z}$ | 7.5 | (Lu *et al.*, 2013) |
| $\lambda_{S,\mu_{200}}$ | 0.1 | (Lu *et al.*, 2013) |
| $\lambda_{S,m_Z}$ | 10 | (Lu *et al.*, 2013) |
| $k_{\mu_{200}}(\mathrm{Hour}^{-1})$ | 0.05 | (Lu *et al.*, 2013) |
| $k_{m_Z}(\mathrm{Hour}^{-1})$ | 0.5 | (Lu *et al.*, 2013) |
| $k_Z(\mathrm{Hour}^{-1})$ | 0.1 | (Lu *et al.*, 2013) |
| $g_Z(\mathrm{Hour}^{-1})$ | 0.1K | (Lu *et al.*, 2013) |
| $S^0_{\mu_{200}}(\mathrm{Molecules})$ | 180K | (Lu *et al.*, 2013) |
| $S^0_{m_Z}(\mathrm{Molecules})$ | 180K | (Lu *et al.*, 2013) |
| $\mu^0_{200}(\mathrm{Molecules})$ | 10K | (Lu *et al.*, 2013) |
| $g_S$ | 18000 | (Lu *et al.*, 2013) |
| $k_S$ | 0.125 | (Lu *et al.*, 2013) |
| $g_{Sl}$ | 50000 | Estimated |
| $k_{Sl}$ | 0.1155 | (Molina-Ortiz *et al.*, 2012) |
| $g_{m_{Sl}}$ | 90 | Estimated |
| $k_{m_{Sl}}$ | 0.5 | Estimated |
| $\lambda_{Sl,\mu_{200}}$ | 0.4 | (Y. N. Liu *et al.*, 2013) |
| $\lambda_{Sl,S}$ | 0.5 | (Nakamura *et al.*, 2018) |
| $\lambda_{S,S}$ | 0.4 | (Peiró *et al.*, 2006) |
| $\lambda_{S,m_{Sl}}$ | 0.5 | (Nakamura *et al.*, 2018) |
| $n_{Sl,\mu_{200}}$ | 1 | (Y. N. Liu *et al.*, 2013) |
| $n_{Sl,S}$ | 3 | (Chen and Gridley, 2013) |
| $n_{S,S}$ | 5 | (Chen and Gridley, 2013) |
| $n_{S,m_{Sl}}$ | 1 | (Chen and Gridley, 2013) |
| $Sl^0_{\mu_{200}}$ | 220000 | Estimated |
| $Sl^0_S$ | 225000 | Estimated |
| $S^0_S$ | 300000 | Estimated |
| $S^0_{m_{Sl}}$ | 180000 | Estimated |
| nIs | 2 | (Jolly *et al.*, 2017) |
| $I^0S$ | 100000 | (Jolly *et al.*, 2017) |
| $\lambda_{I,S}$ | 3 | (Jolly *et al.*, 2017) |
| $g_K$ | 50000 | |
| $k_K$ | 0.1732 | (Gamper *et al.*, 2012) |
| $\lambda_{K,S}$ | 0.5 | (Yori *et al.*, 2011) |
| $\lambda_{K,m_{Sl}}$ | 0.25 | (Y.-N. Liu *et al.*, 2012) |
| $\lambda_{S,K}$ | 0.25 | (Li *et al.*, 2018) |
| $\lambda_{Sl,K}$ | 0.5 | (Y.-N. Liu *et al.*, 2012) |
| $n_{K,S}$ | 2 | (Yori *et al.*, 2011) |
| $n_{K,m_{Sl}}$ | 2 | (Y.-N. Liu *et al.*, 2012) |
| $n_{S,K}$ | 2 | estimated |
| $n_{Sl,K}$ | 4 | (Y.-N. Liu *et al.*, 2012) |
| $\lambda_{K,K}$ | 2 | (Dang, 2002) |
| $n_{K,K}$ | 3 | (Mahatan *et al.*, 1999) |
| $S^0_K$ | 180000 | Estimated |
| $K^0_{m_{Sl}}$ | 300000 | Estimated |
| $Sl^0_K$ | 225000 | Estimated |
| $K^0_K$ | 250000 | Estimated |
| $K^0_S$ | 275000 | Estimated |

**Datasets Used in Kaplan-Meier analysis:**

| Dataset | n(High) | n(Low) |
|---|---|---|
| GSE42568 (Breast Cancer Sample) | 52 | 52 |
| GSE45255 (Breast Cancer Sample) | 67 | 67 |
| GSE3494 (Breast Cancer Sample) | 117 | 119 |
| GSE16445 (Breast Cancer Sample) | 113 | 113 |
| GSE20685 (Breast Cancer Sample) | 163 | 164 |
| GSE20711 (Breast Cancer Sample) | 44 | 44 |
| GSE1456 (Breast Cancer Sample) | 70 | 80 |
| GSE7390 (Breast Cancer Sample) | 99 | 99 |
| GSE37946 (Breast Cancer Sample) | 20 | 20 |
| GSE14814 (Lung Cancer Sample) | 45 | 44 |
| GSE30219 (Lung Cancer Sample) | 147 | 146 |
| CaArray (Lung Cancer Sample) | 234 | 234 |
| GSE14814 (Lung Cancer Sample) | 113 | 113 |
| GSE31908 (Lung Cancer Sample) | 10 | 10 |
| GSE29013 (Lung Cancer Sample) | 27 | 28 |
| GSE19188 (Lung Cancer Sample) | 41 | 41 |
| GSE3141 (Lung Cancer Sample) | 55 | 56 |
| TCGA-LUAD (Lung Cancer Sample) | 37 | 37 |
| GSE31210 (Lung Cancer Sample) | 113 | 113 |
| GSE4573 (Lung Cancer Sample) | 65 | 65 |
| GSE50081 (Lung Cancer Sample) | 91 | 90 |
| GSE37745 (Lung Cancer Sample) | 98 | 98 |
| GSE27651 (Ovarian Cancer Sample) | 30 | 9 |
| GSE30161 (Ovarian Cancer Sample) | 28 | 30 |
| GSE3149 (Ovarian Cancer Sample) | 67 | 49 |
| GSE26712 (Ovarian Cancer Sample) | 60 | 124 |
| GSE15622 (Ovarian Cancer Sample) | 14 | 21 |
| GSE19829 (Ovarian Cancer Sample) | 14 | 14 |
| GSE63885 (Ovarian Cancer Sample) | 40 | 35 |
| GSE18520 (Ovarian Cancer Sample) | 25 | 28 |
| GSE23554 (Ovarian Cancer Sample) | 22 | 6 |
| GSE14764 (Ovarian Cancer Sample) | 37 | 43 |
| GSE26193 (Ovarian Cancer Sample) | 62 | 45 |

**Supplementary figures:**

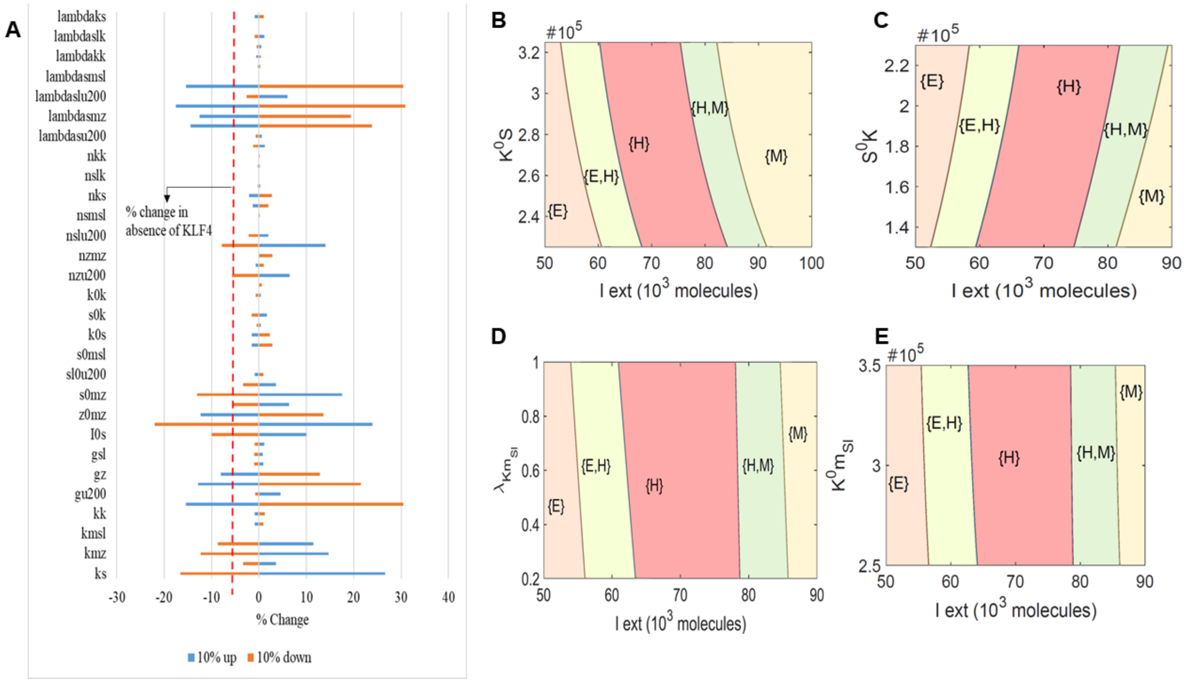


***Fig S1: A)*** Sensitivity analysis indicating percent change in the interval of external EMT signal levels for stable hybrid E/M region, when corresponding parameter values are varied by ±10%. The red dotted line indicates the percent change in the stable hybrid region in the absence of KLF4 (core network) when compared to the coupled network with KLF4. ***B)*** Phase diagrams for the KLF4 network driven by an external signal (I) for varying threshold levels of KLF4 needed for repression on SNAIL. ***C)*** Phase diagrams for the KLF4 network driven by an external signal (I) for varying threshold levels of SNAIL needed for repression on KLF4. ***D)*** Phase diagrams for the KLF4 network driven by an external signal (I) for varying strength of repression on SLUG mRNA by KLF4. ***E)*** Phase diagrams for the KLF4 network driven by an external signal (I) for varying threshold levels of KLF4 needed for repression on SLUG.

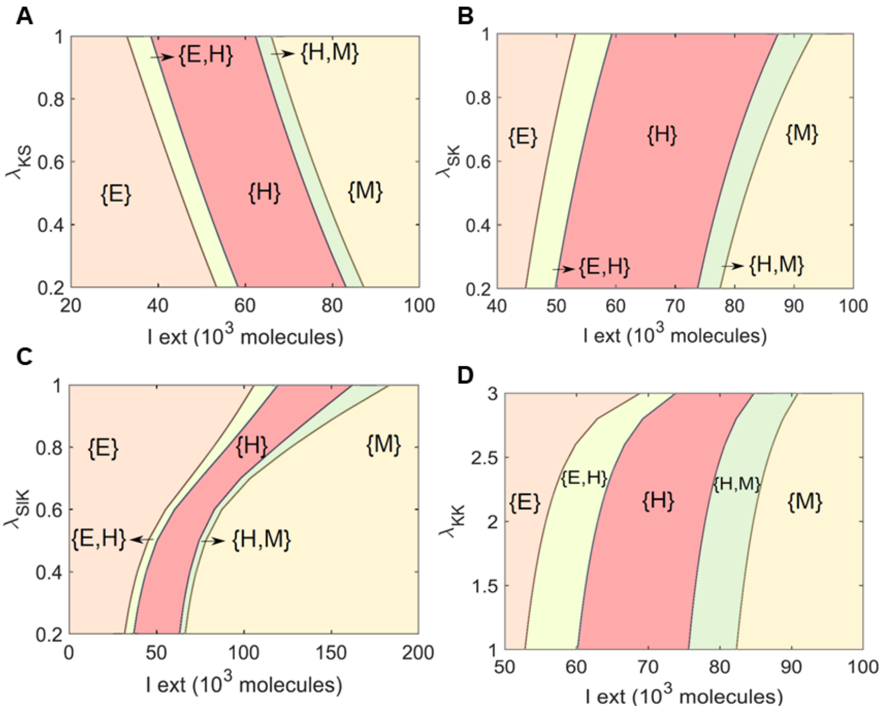


***Fig S2: Effect of SLUG self-activation A)*** Phase diagrams for the KLF4 network with SLUG self-activation driven by an external signal (I) for varying strength of repression on SNAIL by KLF4. ***B)*** Phase diagrams for the KLF4 network with SLUG self-activation driven by an external signal (I) for varying strength of repression on KLF4 by SNAIL. ***C)*** Phase diagrams for the KLF4 network with SLUG self-activation driven by an external signal (I) for varying strength of repression on SLUG by KLF4. ***D)*** Phase diagrams for the KLF4 network with SLUG self-activation driven by an external signal (I) for varying strength of KLF4 self-activation.

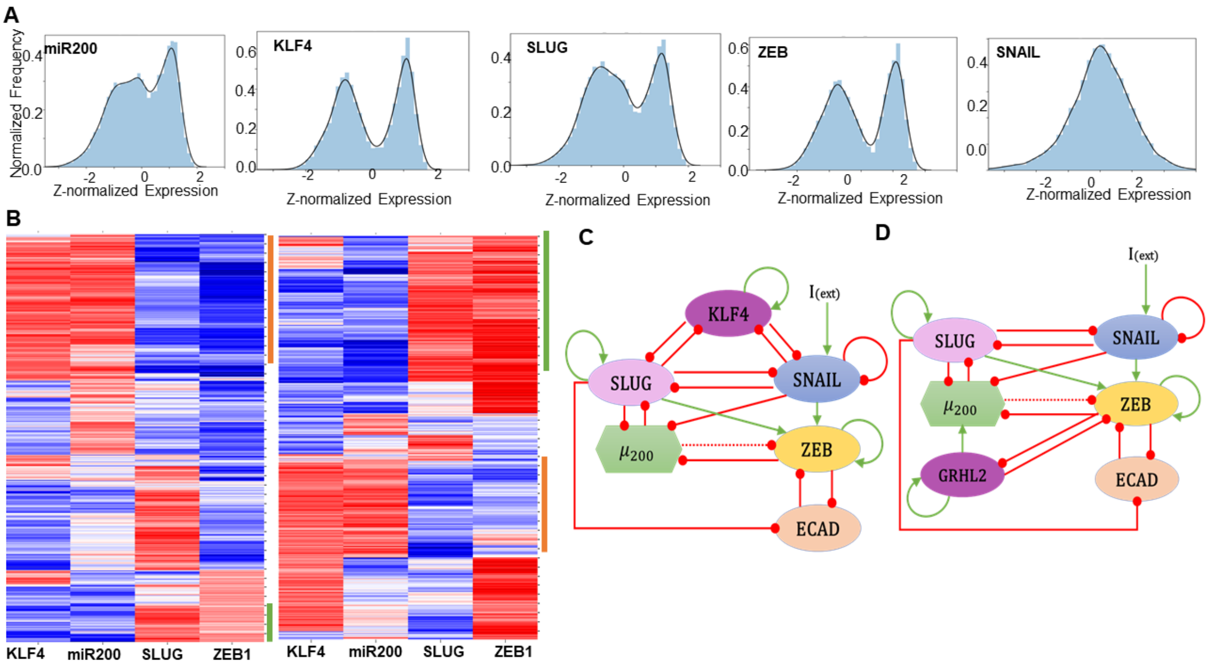

***Fig S3: RACIPE analysis of KLF4-EMT circuit. A)*** Normalized histograms of the expression levels of the nodes in the GRN. ***B)*** Heatmap showing the steady-state expression levels of a single replicate involving ZEB down-expression (ZEB DE) on the simulated GRN ***(Left panel)***. Heatmap showing the steady-state expression levels of a single replicate involving ZEB over-expression (ZEB OE) on the simulated GRN ***(Right panel)***. Green represents mesenchymal phenotype and orange represents epithelial phenotype. ***C)*** Extended KLF4 gene regulatory network ***D)*** GRN containing GRHL2 used for RACIPE simulations

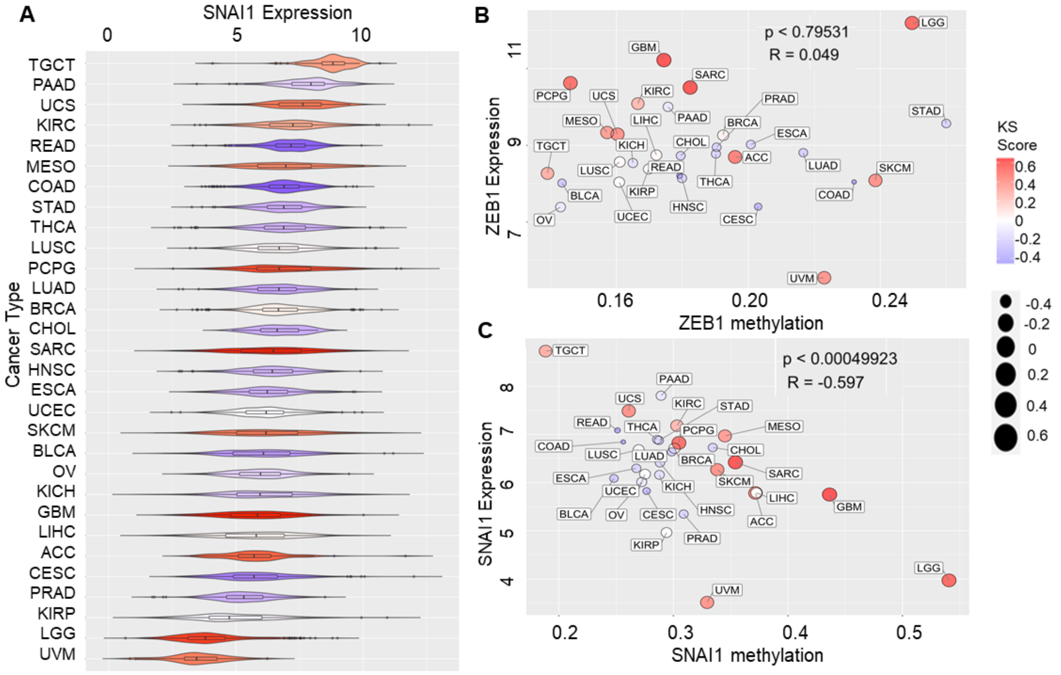


***Fig S4: A)*** SNAI1 expression in TCGA cancers in relation to KS score ***B)*** ZEB1 expression in TCGA cancers in relation to its methylation status ***C)*** SNAI1 expression in TCGA cancers in relation to its methylation status
